# A novel, N_2_-fixing cyanobacterium present and active in the global oceans

**DOI:** 10.1101/2023.08.27.555023

**Authors:** Catie S. Cleveland, Kendra A. Turk-Kubo, Yiming Zhao, Jonathan P. Zehr, Eric A. Webb

**Affiliations:** Marine and Environmental Biology, University of Southern California, Los Angeles, CA, USA; Ocean Sciences Department, University of California, Santa Cruz, Santa Cruz, CA, USA

## Abstract

Marine N_2_-fixing cyanobacteria, including the unicellular genus *Crocosphaera*, are considered keystone species in marine food webs. *Crocosphaera* are globally distributed and provide new sources of nitrogen (N) and carbon (C), which fuel oligotrophic microbial communities and upper trophic levels. Despite their ecosystem importance, only one species, *Crocosphaera watsonii*, has ever been identified and characterized as widespread in the oligotrophic oceans. Herein, we present a novel species, candidatus *Crocosphaera waterburyi* (*C. waterburyi* hereafter), enriched from the North Pacific Ocean, active *in situ*, and globally distributed in environmental datasets. *C. waterburyi* is morphologically, phylogenetically, and physiologically distinct from *C. watsonii*; therefore, description of this novel species provides a new window into previously uncharacterized diversity and ecology of unicellular N_2_-fixing cyanobacterial taxa and further highlights their importance in the global N cycle.

## Introduction

N_2_-fixing cyanobacteria are widespread members of the global oceans and are impactful on the overall health and function of marine ecosystems.^1,2^ Members of the unicellular cyanobacterial genus *Crocosphaera* are photosynthetic, green and orange-pigmented (phycocyanin and phycoerythrin-rich, respectively) bacteria that convert N_2_ gas from the atmosphere into bioavailable forms using the enzyme nitrogenase encoded by the genes *nifH*, *nifD*, and *nifK*^2–4^. Currently, *Crocosphaera* have been described from various biogeographical regions including coastal waters and the oligotrophic oceans^4–6^. The colors of various *Crocosphaera* are indicative of their ecological niches, with the phycocyanin-rich species harvesting red light common in coastal habitats and phycoerythrin-rich strains harvesting blue light available in oligotrophic ocean waters^7^. The coastal, phycocyanin-rich *Crocosphaera* species include: *Crocosphaera subtropica*, *Crocosphaera chwakensis*, and *Cyanothece* sp. BG0011, while the phycoerythrin-rich *Crocosphaera* currently include only one known species, *Crocosphaera watsonii*, which is (previously) the only known abundant, unicellular, free-living, N_2_-fixing cyanobacteria in the oligotrophic oceans^2,5,6^.

*C. watsonii* generates bioavailable nitrogen (N) and carbon (C) and impacts biogeochemical cycling in broad regions^2,4,5^. New C from *Crocosphaera* can provide a resource for upper trophic levels and allows for microbial recycling processes to take place, while new N fuels N-limited phytoplankton that drive the atmospheric biological C pump^2,8^. During summer in the upper euphotic zone of the North Pacific Subtropical Gyre, *C. watsonii nifH* gene-based abundances can often be found at higher cellular abundance than other diazotrophs at 9.4 ± 0.7 × 10^5^ to 2.8 ± 0.9 × 10^6^ *nifH* copies per L^9^. Recent work has also shown that *Crocosphaera* can also have both direct and indirect impacts on N + C export to the deep ocean^10,11^. Deep C export is a mitigating factor in ocean response to rising anthropogenic CO_2_ conditions. Thus, defining the role that *Crocosphaera* plays in both production and export will improve understanding of how the oligotrophic oceans will be impacted by climate change.

In this study, we present the discovery and characterization of a novel, oligotrophic species within genus *Crocosphaera*, Candidatus *Crocosphaera waterburyi* (henceforth, *C. waterburyi*). The *C. waterburyi* Alani8 enrichment was obtained from oligotrophic waters in the North Pacific Ocean near Hawaii. Environmental *nifH* datasets show that *C. waterburyi* is globally distributed in multiple oceans and co-occurs with *C. watsonii* in sinking particle export in sediment traps from 150 to 4,000 m depth. Based on morphological characterization of the enrichment culture, *C. waterburyi* is rod-shaped, ∼5 µm in length by ∼2 µm wide, phycoerythrin-rich, and form large cellular aggregates. The assembled genome of *C. waterburyi* (99% complete, 0% contamination) is comparable in size (5,476,860 bp) and GC content (38.1%) with *C. watsonii* strains (∼37%), yet clusters in a distinct clade when compared phylogenomically, pangenomically, and by average nucleotide identity (ANI). Our characterization of *C. waterburyi* demonstrates it as a previously overlooked, ecologically relevant taxa in oligotrophic ocean regions.

## Materials and Methods

### Cultivation

A single isolate of *C. waterburyi* strain Alani8 was enriched during the 2010 10-day R/V Kilo Moana KM-1013 cruise near Station ALOHA (22° 45′N, 158° 00′W)^12,13^. The enrichment was started from a single, hand-picked *Trichodesmium* colony and incubated in YBCII media without vitamins^14^ at 26°C in a Percival Incubator (Percival Scientific Inc., Perry, IA, USA; 12:12 Light:Dark cycle at ∼100 µmolQ m^-2^ s^-1^). After about 30 days, the *Trichodesmium* colony had lysed, and the culture began to turn orange, suggesting the presence of a phycoerythrin-rich cyanobacterium. Wet mount epi-fluorescent microscopy with Zeiss DAPI and Cy3 filters, a Zeiss AxioStar microscope, and a Zeiss HBO50 light source (Zeiss, Oberkochen, Germany) was used to describe the cellular morphology. Samples from these enrichments were concentrated and streaked on 1.5% Type VII agarose plates (Sigma-Aldrich, Burlington, MA,) and incubated as above for >30 days. This process was repeated twice, and single colonies were picked to obtain unialgal enrichments. Cultures for physiological experiments were non-axenic and were maintained in maximum log growth via weekly transfers to keep heterotrophs in low abundance based on previous *Crocosphaera* culturing work^5^.

### Growth and Nitrogen Fixation Rates: *C. waterburyi* and *C. watsonii*

N_2_ fixation rates of *C. waterburyi* Alani8 and *C. watsonii* WH0003 were measured in triplicate using the acetylene reduction assay^15^. Triplicate Nalgene 43 mL polycarbonate tubes contained 30 mL of each culture and 13 mL of headspace. The tubes were sealed with tightened septa caps (I-CHEM, Calhoun, LA, USA), charged with >10% acetylene synthesized from carbide (Thermo Fisher Scientific Inc., Waltham, MA, USA), and produced ethylene was measured on a GC-8a gas chromatograph (Shimadzu, Kyoto, Japan) following 12 hours incubation. Acetylene reduction was calculated using 9 ppm ethylene standards (Cambridge Isotopes, Cambridge, MA, USA). Final rates were normalized to raw in-vivo chlorophyll a fluorescence due to difficulty of cell counts within aggregates.

Growth rates of *C. watsonii* and *C. waterburyi* were determined from triplicate in-vivo chlorophyll fluorescence data obtained on a TD-700 Fluorometer (Turner Designs, Carlsbad, CA, USA). Cultures were grown under the following conditions: cool white lights, ∼150 µmol Q m^-2^ s^-1^, 12:12 diel cycle, and 26°C. Before each measurement, cultures were thoroughly mixed to fully homogenize aggregates in solution. Growth rates were calculated using the raw fluorescence increase and the Malthusian growth model in GraphPad Prism (La Jolla, CA, USA). The temperature profile of *C. waterburyi* was generated using the same methods but under slightly different light conditions (3,000 K warm white light and 96 µmol Q m^-2^ s^-1^ light intensity). All lights for the temperature experiment were identical, and Percival incubators were used for all growth curves.

### Extraction and Sequencing

To concentrate biomass for DNA extraction, 100 mL of culture was centrifuged at 13,000 RPM for 2 minutes at 25°C to form a pellet. DNA was then extracted using the Qiagen DNeasy PowerBiofilm kit (Qiagen, Germantown, MD, USA) following the manufacturer’s protocol with the following modifications: after addition of the cell material to the bead beating tube, the cells were lysed with liquid N_2_ freeze-thaws (5X), tube agitation (3X), and 65°C overnight Proteinase K (20 ng/µL; VWR International, Radnor, PA, USA) incubation. DNA was quantified using a Qubit 4 fluorometer (ThermoScientific, Waltham, MA, USA), and 260/280 quality was verified with a NanoDrop 1,000 spectrophotometer (Thermo Fisher Scientific, Waltham, MA, USA). Library preparation with the NEBNext® DNA Library Prep Kit and PE150 sequencing at a depth of 1Gbp was completed at Novogene Inc. (Sacramento, CA, USA).

### Genome Assembly

The reads were assembled on the open-source web page KBase (KBase.com) following the public narrative, “Genome Extraction for Shotgun Metagenomic Sequence Data.” The assembly pipeline is briefly summarized as follows: FastQC v0.11.9 checked read quality^16^, Trimmomatic v0.36 removed contaminating and low scoring sequences with a sliding window size of 4 and minimum quality of 15 (0.1% reads dropped)^17^, Kaiju Taxonomic Classifier v1.5.0 described the taxonomic distribution of the reads^18^, MetaSPAdes v3.13.0 assembled the reads with default parameters^19^, MaxBin2 v2.2.4 binned the contigs to obtain metagenome assembled genomes (MAGs)^20^, CheckM v1.0.18 analyzed quality^21^, and GTDB-tk v1.7.0 determined the taxonomy of *C. waterburyi.*^22^

### Phylogenetic Tree Construction

To place the *C. waterburyi* genome in context with other near relative genomes available in GenBank, accessions of family *Aphanothecacea* and genus *Cyanothece* were obtained from the NCBI assembly site. An initial phylogenomic tree with 257 genomes/MAGs was created using the GToTree workflow and associated programs^23–28^ with *Gloeobacter violaceus* PCC 7421 (GCA_000011385.1) as the root. Subsequently, another tree was created using only the 35 assemblies most closely related to *C. waterburyi*. The tree used 251 conserved cyanobacteria HMMs^27^ with at least 50% of the HMMs required in each genome to be included in the tree. The output tree data from GToTree was piped into IQTree2 using the best model finder method and 1,000 bootstraps^29,30^ to generate the final consensus tree.

We additionally used NCBI-blastn to place the *C. waterburyi nifH* gene in an environmental context. A phylogenetic tree was created using the *nifH* gene sequences from *Crocosphaera* enrichment cultures and 250 *nifH* gene sequences identified by blastn as having high average nucleotide identity to the *C. waterburyi nifH* gene (**Suppl. table S3**). These combined sequences were aligned in Geneious^31^ using Clustal Omega 1.2.2^32^, trimmed, and a subsequent *nifH* gene tree was created using RAxML 8.2.11^33^ with a GTR GAMMA nucleotide model, rapid bootstrapping (1,000 bootstraps), and the maximum likelihood tree algorithm.

### Pangenome Analysis

We used the pan genomic pipeline in Anvi’o v7.1^34^ to define the core and accessory genomes of 10 *Crocosphaera* assemblies, including six *C. watsonii* strains (WH0003 (GCA_000235665.2), WH0005 (GCA_001050835.1), WH0402 (GCA_001039635.1), WH8501 (GCA_000167195.1), WH8502 (GCA_001039555.1), WH0401 (GCA_001039615.1)), *C. chwakensis* CCY0110 (GCA_000169335.1), *C. subtropica* ATCC 51142 (GCA_000017845.1), *Cyanothece* sp. BG0011 (GCA_003013815.1), and the novel *C. waterburyi*. Two environmental MAGs, *Crocosphaera* sp. DT_26 (GCA_013215395.1) and *Crocosphaera* sp. ALOHA_ZT_9 (GCA_022448125.1), were excluded from the pangenome as they were not from isolated cultures^35,36^ and their physiology has not yet been characterized. All assemblies, beside *C. waterburyi*, were obtained from NCBI. Briefly, the genomes were reformatted and annotated with NCBI-COGS, Pfams, KOfams, and HMMER^27,37–40^ to define the conserved gene content in each assembly. The pangenome was constructed using an MCL 2 threshold suitable for less-similar genomes, and the FastANI^41^ heatmap used an ANI lower threshold of 80% similarity (**Suppl. table S2**). Genomes were ordered by ANI similarity, and gene clusters were aligned and ordered in Anvi’o by presence or absence in the genomes.

### Environmental Read-Mapping

We used the *C. waterburyi*, *C. watsonii* WH0003, *C. chwakensis* CCY0110, *Cyanothece* sp. BG0011, and *C. subtropica* ATCC 51142 genomes as targets for read recruiting to sixty-three metagenome samples from 4,000 m depth in the ALOHA Deep Trap Sequencing project (PRJNA482655; DeLong research group at University of Hawai’i and Simons Collaboration on Ocean Processes and Ecology)^35^, GoShip surface metagenomes^42^, TaraOceans metagenomes^43^, and BioGEOTRACES metagenomes^44^. Briefly, the pipeline was as follows: Bowtie2 mapped reads to the contig set^45^, Samtools v1.9 converted SAMs to BAMs^46^, CoverM filtered the BAMs at 98% identity over at least 70% of the read (https://github.com/wwood/CoverM), and Anvi’o v7.1 visualized and parsed the results^34^. Genome presence or absence across samples was determined using a threshold of ≥25% genome detection as per previous methodology^47^ and the Anvi’o % recruitment metric (interpreted as: “of the reads that were mapped, X% of them mapped to genome A and X% mapped to genome B) was used as a metric of relative abundance^34^.

### Detection of *nifH* Gene and Transcripts in the North Pacific Subtropical Gyre

Samples for the determination of diazotroph community composition and activity were collected during the SCOPE-PARAGON I research expedition in the North Pacific Subtropical Gyre (NPSG) July 22-August 5, 2022 (R/V Kilo Moana). Three types of samples were collected: size fractionated seawater samples (DNA); diel seawater samples (RNA); and samples of particles sinking out of the euphotic zone (DNA/RNA). All seawater samples were collected from three depths, 25 meters, 150 meters, and the deep chlorophyll maximum (DCM: ∼135 meters), using Niskin® bottles mounted to a CTD rosette (SeaBird Scientific Bellevue, WA, USA), and transferred into acid-washed polycarbonate bottles or carboys. Large volume (20 L) seawater samples were filtered serially using gentle peristaltic pumping through the following filters: 100 µm nitex mesh (25 mm, MilliporeSigma, Burlington, MA, USA); 20 µm polycarbonate (25 mm; Sterlitech Corp., Auburn, WA, USA) 3.0 µm polyester (25 mm, Sterlitech Corp., Auburn, WA, USA); and 0.2 µm Supor® (25 mm; Pall Corporation, Port Washington, NY, USA). Diel samples (2.5-4 L) were collected every ∼6 hr over 30h and filtered serially through 3.0 µm polyester (25 mm, Sterlitech Corp., Auburn, WA, USA) and 0.2 µm Supor® filters (25 mm; Pall Corporation, Port Washington, NY, USA), with care taken to keep filtration times under 30 min.

Sinking particles were collected using surface tethered net traps (diameter 1.25 m, 50 µm mesh cod end^48^ and deployed at 150 m for 24 hr. Upon recovery of the net traps, particles were gently resuspended in sterile filtered 150m water and split into multiple samples as previously described^49^. Particles slurries were gently filtered through 0.2-µm Supor® filters (25 mm; Pall Corporation). All filters were flash frozen in liquid N_2_ and stored at −80°C until extraction.

DNA and RNA were co-extracted from all samples using the AllPrep DNA/RNA Micro kit (Qiagen, Germantown, MD, USA) according to the manufacturers’ guidelines with modifications described previously^50^. RNA extracts were DNase digested using the Turbo DNA-free kit (Ambion, Austin, TX, USA) to remove any DNA contamination, then cDNA was synthesized with the Superscript IV First-Strand Synthesis System (Invitrogen, Waltham, MA, USA) primed by universal *nifH* reverse primers nifH2, nifH3 using reaction conditions as previously described^51^. All DNA and RNA extracts were screened for purity using a NanoDrop spectrophotomer (ThermoScientific, Waltham, MA, USA), and DNA was quantified using Picogreen® dsDNA Quantitation kit (Molecular Probes, Eugene, OR, USA).

Partial *nifH* fragments were PCR-amplified using the universal primers nifH1-4^52,53^ and sequenced using high throughput amplicon sequencing as detailed previously^54^. Amplicon sequence variants (ASVs) were defined using the DADA2 pipeline^55^ with customizations specific to the *nifH* gene (J. Magasin, https://github.com/jdmagasin/nifH_amplicons_DADA2). *Crocosphaera* ASVs were identified using blastx against a curated *nifH* genome database (www.zehr.pmc.ucsc.edu/Genome879/), including ASVs 100% identical to *C. waterburyi* and *C. watsonii* WH8501 (AADV02000024.1). Demultiplexed raw sequences are available under BioProject PRJNA1009239 in the Sequence Read Archive at NCBI.

## Results and Discussion

### Morphological and Physiological Characteristics

#### a. *C. waterburyi* displays morphological similarities with both coastal and oligotrophic *Crocosphaera*

Following isolation from the North Pacific near Station ALOHA, it was observed that *C. waterburyi* consistently displays cell morphology and pigmentation that bridges the evolutionary gap between the coastal, phycocyanin-rich *C. subtropica*, *C. chwakensis*, and *Cyanothece* sp. BG0011 (CrocoG hereafter) with the oligotrophic, phycoerythrin-rich *C. watsonii*. Specifically, *C. waterburyi* cells are rod-shaped and ∼5 µm long by ∼2 µm wide (**Figure 1A**) like the coastal *Cyanothece* sp. BG0011,^56^ yet are phycoerythrin-rich like the spherical *C. watsonii* (**Figure 1B**). *C. waterburyi* also forms aggregates in culture (i.e., flocs) similar to the coastal *Crocosphaera* species^6^. However, *C. waterburyi*-like cells also appear to be present sympatrically with *C. watsonii*-like cells in particle export traps from the North Pacific Ocean over multiple years (F**igure 1C-E**).

**Figure 1.**
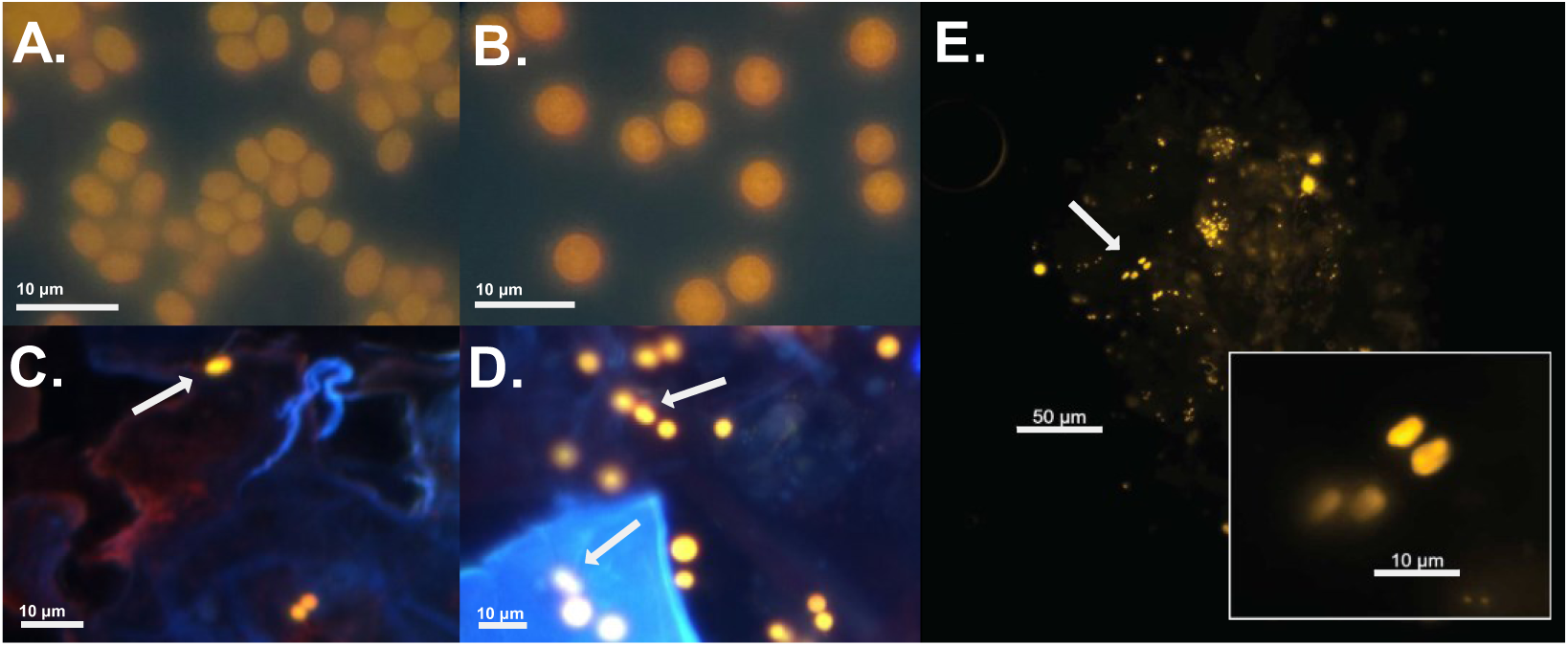
Morphological difference between **(A)** and *C. watsonii* **(B)** as visualized by an epifluorescent microscope with a DAPI LP excitation/emission. Environmental photos taken using DAPI Long Pass excitation from 75 m depth net traps **(C, D)** show a mixture of spherical and rod-shaped (indicated by white arrows) cells during the 2010 North Pacific RV Kilo Moana KM1013 cruise from which *C. waterburyi* was isolated. *C. waterburyi*-like cells (i.e., rod-shaped, phycoerythrin-rich) visualized by a Cy3 filter were also attached to sinking particles caught in net traps during the 2022 SCOPE-PARAGON I research expedition **(E)**.

**Figure 2.**
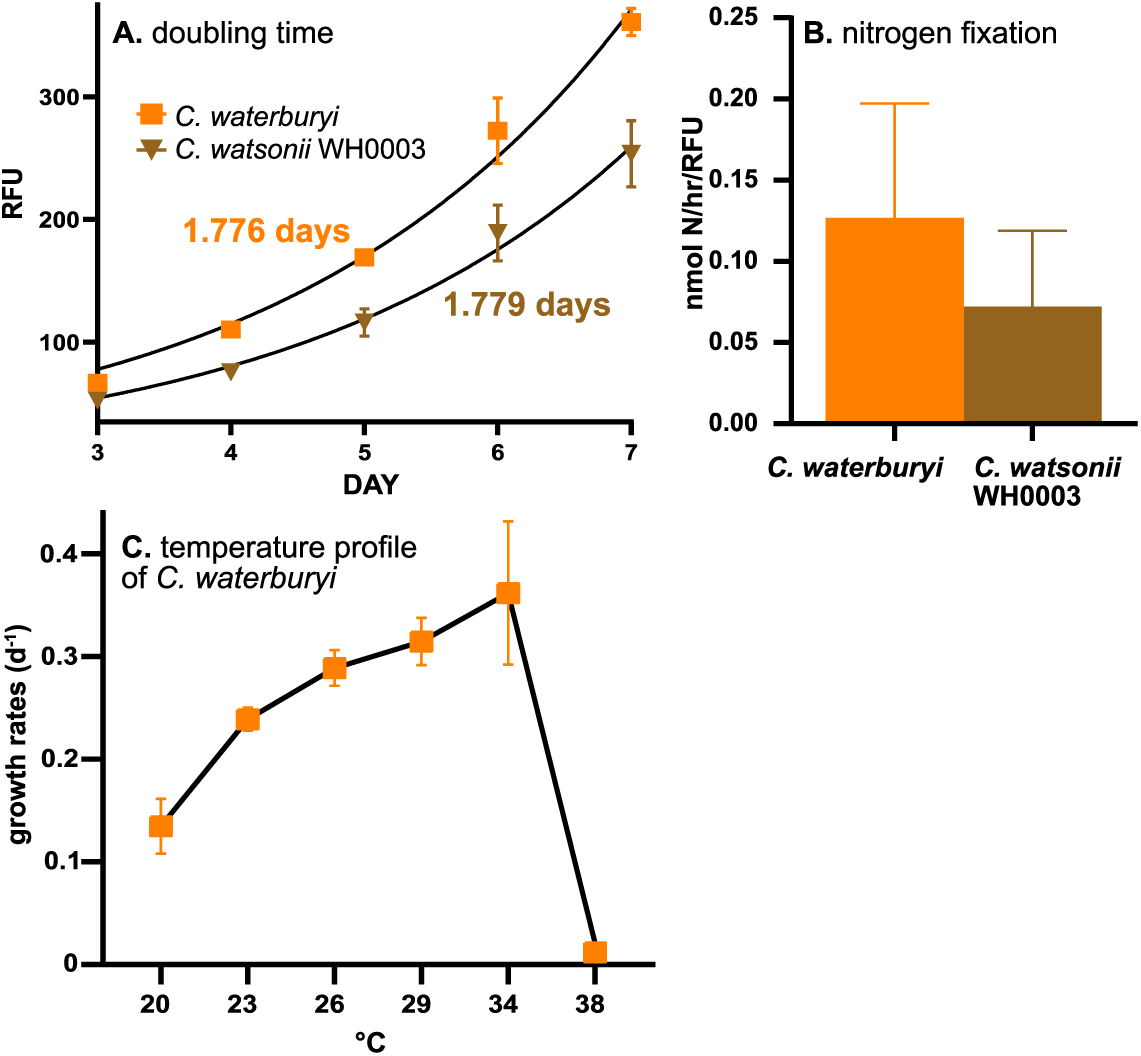
Growth **(A)** and N_2_ fixation rates **(B)** of representative oceanic *Crocosphaera* species and the temperature range of *C. waterburyi* **(C)** at 96 µmol Q m^-2^ s^-1^ and a 12:12 diel cycle.

#### b. *C. waterburyi* may outcompete *C. watsonii* in projected warmer oceans

Growth and N_2_ fixation rates were measured to compare the physiology of *C. waterburyi* to *C. watsonii* and to further understand what role *C. waterburyi* has in N + C cycling in the ocean. When grown under ∼150 µmol Q m^-2^ s^-1^, a 12:12 diel cycle, and 26°C, *C. waterburyi* was found to have a slightly faster doubling time (1.776 vs 1.779) and nitrogen fixation rate (0.127 ± 0.04 nmol N hr^-1^rfu^-1^ vs 0.072 ± 0.03 nmol N hr^-1^rfu^-1^) than *C. watsonii* WH0003 (**Figure 3A-B**), although these differences were not statistically significant.

**Figure 3.**
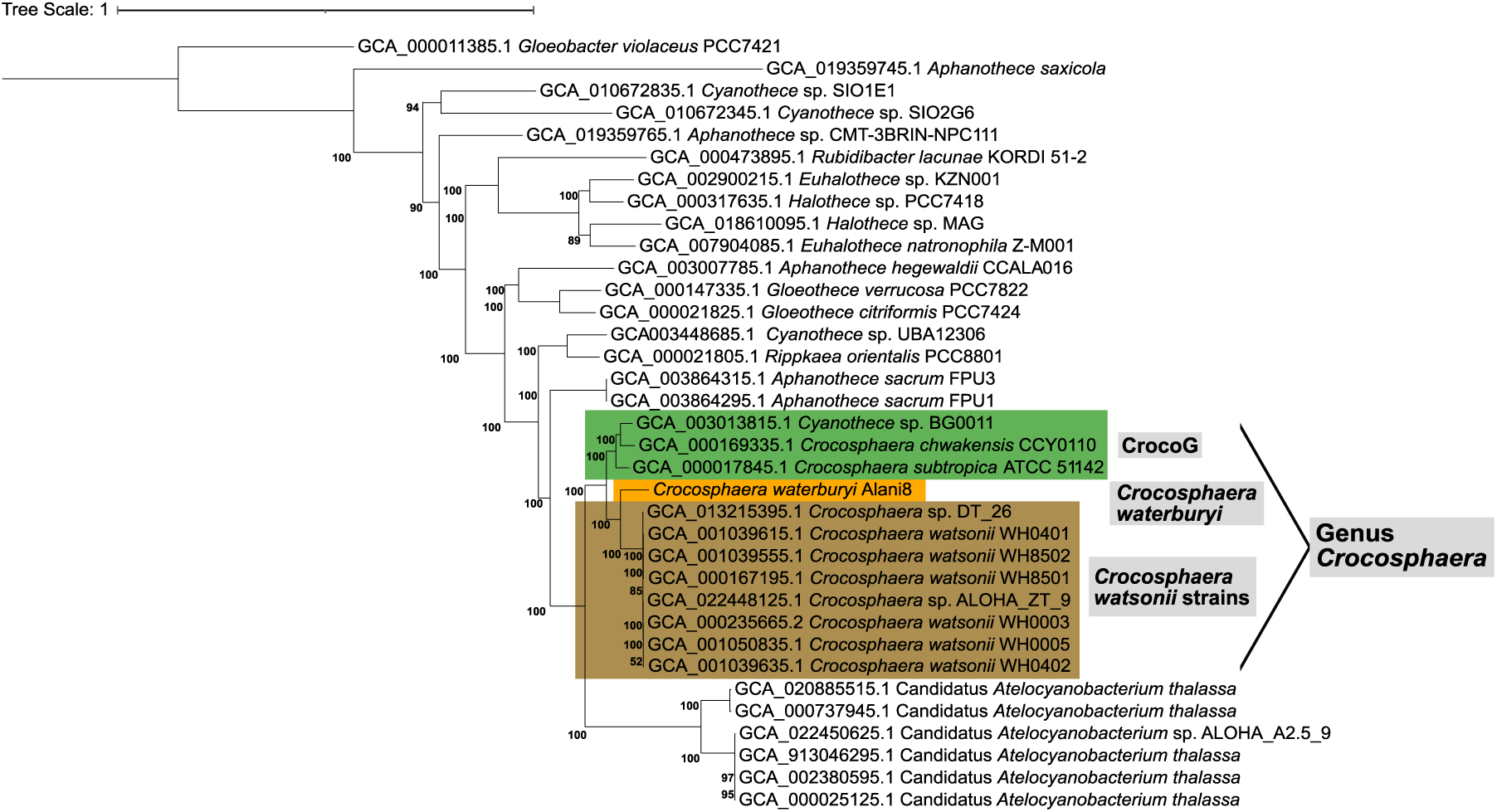
The phylogenomic tree of 35 cyanobacterial taxa from NCBI that are most closely related to *C. waterburyi*. The CrocoG subclade is denoted by green highlighting, *C. watsonii* by brown highlighting, and *C. waterburyi* by orange highlighting. Bootstrap values below 85% are not shown.

However, despite having the same doubling time and N_2_ fixation rate at 26°C, *C. waterburyi* was found to have a broader optimal temperature range than *C. watsonii* WH0003. *C. watsonii* have been recorded to grow optimally from 26-32°C with growth dropping off to ∼0.1 at 23°C and 34°C with death occurring at 35°C^57^. However, under similar conditions, *C. waterburyi* was found to grow faster than *C. watsonii* at both lower (0.24 d^-1^ at 23°C) and higher (0.36 d^-1^ at 34°C) temperatures (**Figure 3C**). *C. waterburyi* was also found to have a broad temperature optimum and maintained growth rates from 26-34°C that were statistically the same (**Figure 3C**). While a colder lower cardinal temperature may allow *C. waterburyi* to grow in higher latitudes than *C. watsonii*, a higher upper cardinal temperature indicates that *C. waterburyi* may also grow better than *C. watsonii* under future climate change-induced ocean warming conditions.

### Evolutionary Relationships

#### a. *C. waterburyi* represents a novel subclade within the genus *Crocosphaera*

An initial phylogenomic tree was created with 257 genomes within the family *Aphanothecacea* and genus *Cyanothece* (curated at NCBI Assembly) to ensure correct taxonomic placement of *C. waterburyi*. Following this, a subsequent tree was made using only the 35 taxa most closely related to *C. waterburyi* (**Figure 3**). Based on this, *C. waterburyi* was found to be most closely related to *C. watsonii* yet still clustered in a separate subclade. *C. watsonii* and *C. waterburyi* do, however, form an ‘oceanic’ group within the genus, which is distinct from the coastal CrocoG (**Figure 3**).

Previously, *C. watsonii* have been found to display strain specific differences in cell size and exopolysaccharide (EPS) production^5,58^. However, despite these differences, the *C. watsonii* strains are all phylogenomically closely related (**Figure 3**). *C. waterburyi* displays both morphological (**Figure 1**) and strong phylogenomic differences from *C. watsonii* (**Figure 3**), further supporting its description as a novel species of *Crocosphaera*.

### Pangenomic Comparisons of Genus *Crocosphaera*

The full genomic potential of the genus *Crocosphaera* has never been characterized, and thus, how gene content varies across the genus and in *C. waterburyi* has never been defined. To ensure that only high quality genomes were included in the *Crocosphaera* pangenome, CheckM^21^ was used to demonstrate that all genomes were >98% complete, <2% contamination with N50 values between 9,214 and 4,934,271 (**Suppl. Table S1**). The draft genome of *C. waterburyi*, specifically, was found to be high quality at 99.56% complete, 0% contamination, and an N50 of 69,427. Overall, all *Crocosphaera* genomes used for subsequent analyses were also found to be larger than many marine bacterial genomes^59^. The GC content of *C. waterburyi* (38.1%) was slightly higher than the *C. watsonii* strains (37.1 - 37.7%) but comparable to the coastal *Cyanothece* sp. BG0011 genome in the CrocoG subclade (38.2%).

#### a. Pangenomics and average nucleotide identities recapitulate the phylogenomic placement of *C. waterburyi*

Genus *Crocosphaera*, despite their wide biogeographical range and habitat difference (coastal vs oligotrophic), have 2,391 gene clusters shared in their “genomic core,” (**Figure 4**). The core genes are enriched in distinct functions related to the lifestyle of these organisms, including N_2_-fixation, phosphate uptake, iron (III) utilization, photosynthesis-related, phycocyanin and phycoerythrin, and mobile genetic element-related genes (**Suppl. Table S4**).

**Figure 4.**
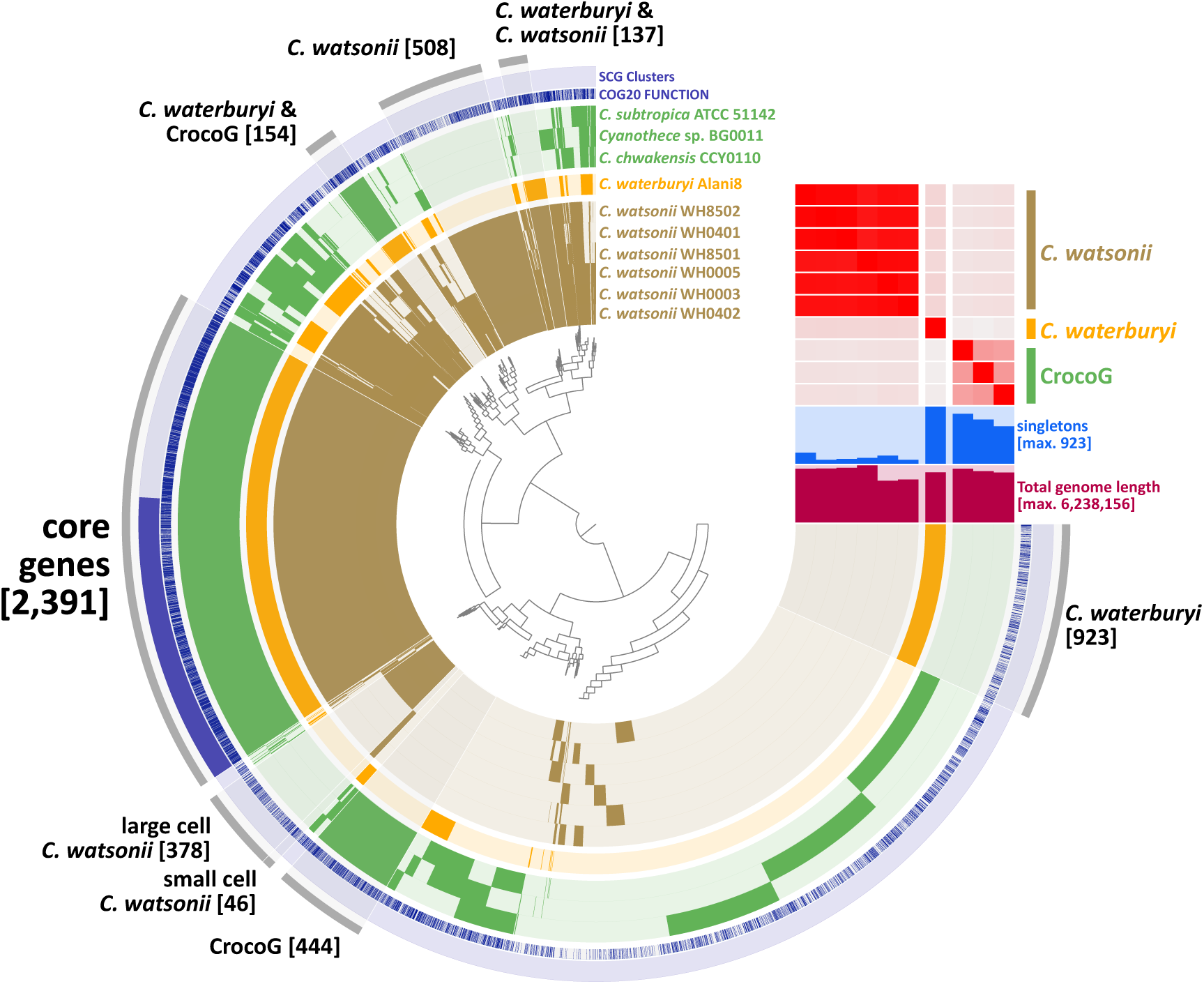
The pangenome of the genus *Crocosphaera*. The heatmap shows % ANI similarity and subclade distinctions of the genus with a lower threshold of 80% similarity, and the tree at the center shows gene cluster presence vs absence. The numbers in brackets indicate the number of gene clusters in each bin. The gene annotations are shown in blue in the “COG20 Function” layer and the single copy genes in all 10 genomes are shown in the “SCG Clusters” layer. The “Singletons” (shown above “Total genome length”) are the number(s) of gene clusters present only in individual genomes.

Pangenomic analysis also revealed that members of each phylogenomically-defined *Crocosphaera* clade have auxiliary genes found only in those groups. For example, CrocoG and *C. watsonii* subclades each had genes distinct to their groups (each sharing 444 and 508 gene clusters, respectively; **Figure 4**), enriched in different mobile genetic element-related genes (**Suppl. Table S4**). *C. watsonii* also showed sub-grouping at the strain level with the small cell phenotypes sharing 46 unique gene clusters and the large cell phenotype sharing 378 gene clusters (**Figure 4**). Overall, *C. waterburyi* was found to have the most unique, auxiliary genes with a total of 986 genes and 923 gene clusters (**Figure 4**), although 51% lacked annotation by NCBI-COGS, Pfam, and KOfam. These high auxiliary gene numbers in *C. waterburyi* could be due to only one genome being available from this group. However, broad groupings based on the presence and absence of genes in core and auxiliary genome do corroborate the phylogenomic structure. *C. waterburyi* was also found to share distinct gene clusters with the CrocoG (154 gene clusters) and separately with *C. watsonii* strains (137 gene clusters), (**Figure 4**), further affirming it as an evolutionary intermediate between the oligotrophic, open ocean *C. watsonii* and the coastal CrocoG.

When further compared by average nucleotide identity (ANI) and visualized in a heatmap (>80% lower threshold), *Crocosphaera* were again differentiated into the same 3 subclades: the *C. watsonii* strains, the CrocoG, and *C. waterburyi*. As expected, the six *C. watsonii* strains had high ANI similarity at >98%. However, *C. waterburyi* was only 82% ANI to all cultured *C. watsonii* strains and 80-81% to the CrocoG (**Suppl. Table S2**). As these values are below both the intra-species 95% ANI cutoff and the 83% ANI inter-species value^41^, they further support that the sympatric *C. watsonii* and *C. waterburyi* do represent different oligotrophic ocean species. In summary, based on both gene content and ANI, *C. waterburyi* represents a transitional taxon that bridges features of the previously described green, coastal, and orange, oligotrophic *Crocosphaera* subclades. As well, it appears to be evolving sympatrically with *C. watsonii* in the oceans.

#### b. Gene contents of *C. waterburyi* and *C. watsonii* reveal functional differences of these sympatric species

While *C. waterburyi* and *C. watsonii* do appear to be similar in core functions (**Figure 4**) and habitat, there are characteristics of each that may impact their role in the ocean. One prime example is the presence of a CRISPR-Cas type I-B system in *C. waterburyi* (**Suppl. Figure S1-S2**) but not in any of the 6 *C. watsonii* strains. *C. watsonii* only encodes Csa3, which is annotated as a transposase and is not a true Cas gene^60^, (**Suppl. Table S4-S5**). CRISPR-cas systems, present in some bacteria, provide them with immunity against bacteriophage infection^61^, and cyanobacteria overall have an abundance of the Type III-B system^62^, including the sympatric cyanobacterium *Trichodesmium thiebautii*^60^. However, based on analysis with CCTyper and Anvi’o, *C. waterburyi* and other closely related single-celled cyanobacteria additionally encode the Type I-B system (**Suppl. Figures S1, S2**). With this I-B CRISPR-cas system, *C. waterburyi* may be more resistant to cyano-phage infection than *C. watsonii*, suggesting a species-level difference in C + N cycle impacts via viral lysis.

Additionally, despite sharing some Fe (III) and (II) utilization genes in the genomic core (*FeoAB, AfuA, FbpB*, **Figure 4**; Core), *C. waterburyi*, *C. watsonii*, and the CrocoG were found to have variation in other predicted Fe acquisition genes. This finding is relevant as Fe demand is increased in oligotrophic ocean diazotrophs relative to other phytoplankton due to their obligatory Fe requirement of the metalloenzyme nitrogenase. Fe can also be very limiting in the environment^63–65^ and Fe limitation has been found to have a strong effect on N_2_-fixation rates in *Crocosphaera*^66–68^. *C. waterburyi* was found to encode additional, unique copies of *feoAB* Fe (II) transport genes (**Figure 4**; **Suppl. Table S4, S6**). Blastp even identified this additional *C. waterburyi feoA* (PDB:2GCX) and *feoB* (PDB:3TAH) as more similar by % ANI to *feoAB* in *Gloeocapsa* sp. PCC 73106 (WP_006528539.1, WP_006528538.1), which are of freshwater origin^69^, highlighting a hereditary difference in this core metabolism and a potential horizontal gene transfer event. As mentioned, Fe availability has been found to be a major limiting nutrient for diazotrophs in the ocean, so it is relevant to note that evolutionary differences in Fe utilization genes in the marine *Crocosphaera* may have impacts on their ability to scavenge it from the oceans.

In summary, *Crocosphaera*, including novel *C. waterburyi*, are overall similar in GC content, genome size, and core metabolic features. However, distinct genetic functions, such as differences in Fe utilization genes and predicted phage immunity, distinguish the oceanic species, *C. watsonii* and *C. waterburyi,* and imply that they have somewhat different ecological roles in the environment.

### Presence and Activity in the Oligotrophic Ocean

The Earth’s warm oligotrophic oceans are characterized as low-nutrient, high microbial remineralization regions, and unlike the coastal ocean, these oceanographic ‘deserts’ are vast in size, comprising >60% of the global oceans^70^. Due to consistently low nutrients, organisms living in these ecosystems rely heavily on N_2_ fixation by diazotrophs, including *Crocosphaera*, in the euphotic zone to fuel microbial to upper trophic level productivity^1,2,8^. Therefore, determining where oligotrophic *Crocosphaera* species are present and active is important for understanding their contributions to global biogeochemistry.

#### a. *C. waterburyi* is present and active in global ocean euphotic zones

*C. watsonii* distribution and abundance has been previously well characterized in the North Pacific Ocean near Station ALOHA,^3,9,11,71^ and they have been observed as consistent members of the bacterial community. However, despite being isolated from the North Pacific Ocean near Station ALOHA and being sympatric with *C. watsonii*, *C. waterburyi’*s *in situ* abundances and activity are previously un-characterized.

To remedy the lack of *C. waterburyi* environmental data, we utilized a summer 2022 *nifH* amplicon DNA/RNA dataset from the Station ALOHA region. This showed that the *C. waterburyi nifH* gene had high relative abundances, particularly in the 20 and 100 µm size fractions, at the DCM and in 150 m depth samples (**Figure 5A**). As *C. waterburyi* cells are only ∼5 µm long (**Figure 1**), their presence in larger size fractions (20 and 100 µm) provides evidence that these cells likely form large aggregates *in situ*, as has been observed in culture with the Alani8 strain (**Figure 1**). Transcripts 100% identical to *C. waterburyi nifH* were detected in the early evening (18:15) in the 3-µm size fraction at 150 m depth, and *C. waterburyi* DNA was found at high % relative abundance at 150 m across multiple days (**Figure 5C**). However, contrastingly, *C. watsonii nifH* transcripts were only found at the DCM (**Figure 5B**), and *C. watsonii* DNA did not have a high % relative abundance in any size fraction across sampling dates (**Figure 5C**). These data suggest that, not only is *C. waterburyi* present in the North Pacific Ocean, but it is transcriptionally active and contributing to C + N export through sinking. Also, despite overlapping in the same larger ocean regions, *C. waterburyi* and *C. watsonii* seemingly display a depth difference in *nifH* transcript presence during this sampling event.

**Figure 5.**
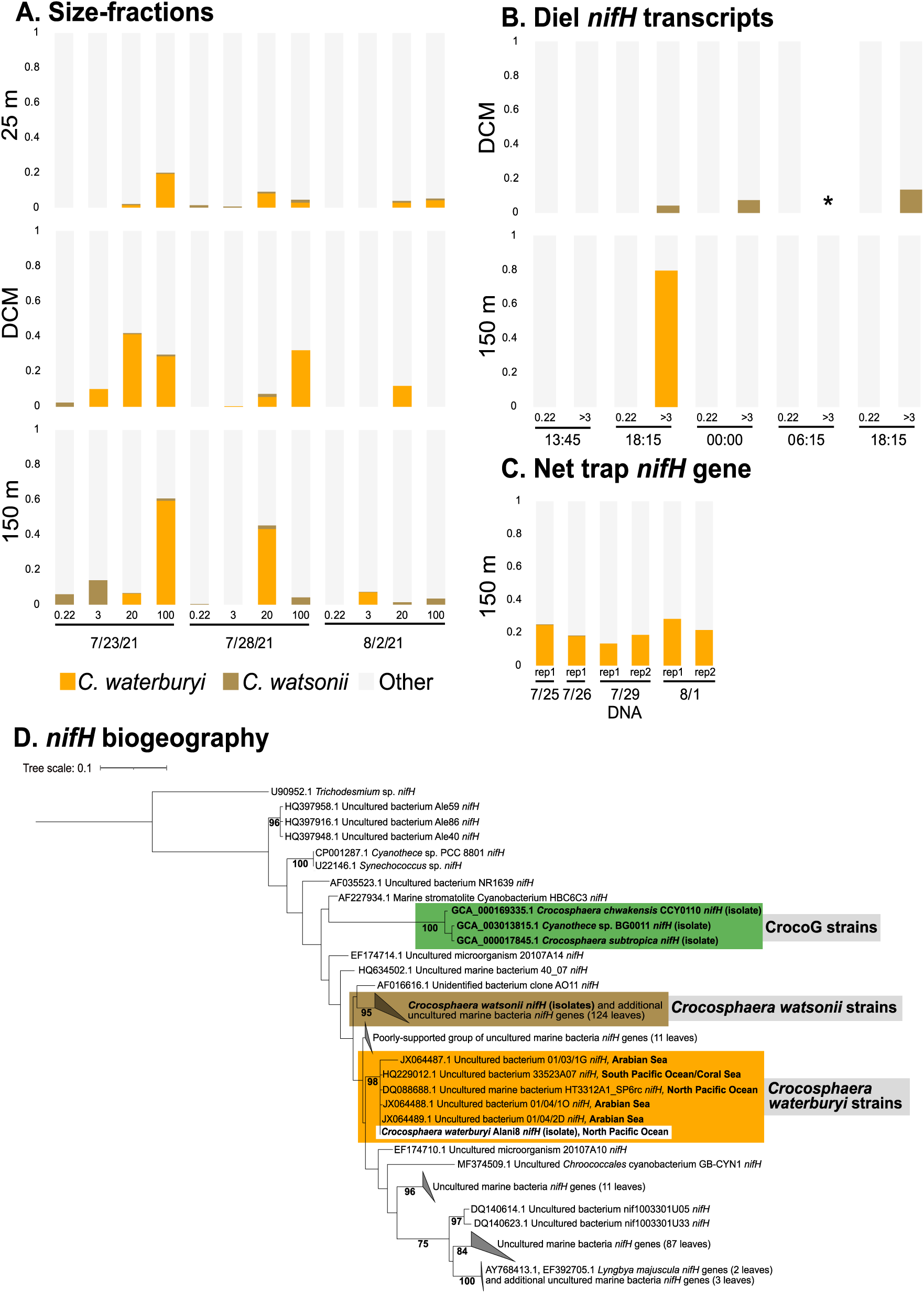
The *nifH* gene relative of abundance of *C. waterburyi*, *C. watsonii*, and other diazotrophs in the North Pacific Ocean. Shown are the size-fractionated *nifH* gene relative abundance from deployed net traps **(A)**, *nifH* transcripts from a diel sampling over the course of a day **(B)**, and the *nifH* gene presence over four days **(C)**. The DCM fell at a depth of 135 m, and data is not available for one DCM >3-µm size fraction sample over the diel sampling (marked with an asterisk “*”). **(D)** shows the *nifH* DNA phylogeny of 250 NCBI-blastn hits closest to *C. waterburyi*.

In addition to the *nifH* amplicon dataset collected from the North Pacific, other *nifH* amplicon datasets were examined using blastn to determine the presence of *C. waterburyi* in other oceans. Our *C. waterburyi* enrichment *nifH* sequence was a 100% ANI matched to a *nifH* amplicon sequenced from the Arabian Sea^72^ (**Figure 5D**). Phylogenetically, the *C. waterburyi nifH* also clustered with additional environmental *C. waterburyi*-like sequences from the North Pacific Ocean, South Pacific Ocean/Coral Sea, and Arabian Sea (**Figure 5D**). Therefore, while this species was originally isolated from and is active in the North Pacific Ocean, its biogeographical range extends much farther. Finally, in addition to the phylogenomic subclades characterized in this study (*i.e.,* CrocoG, *C. watsonii*, and *C. waterburyi*), there are five additional, well-supported *nifH* sub-clades closely related to *C. waterburyi* (**Figure 5D**) with uncultured diversity that are of interest for future isolation efforts.

In the North Pacific, *C. waterburyi* was also recently identified as the novel, uncultured “Croco_otu3,” which is at higher relative abundance deep in the euphotic zone, appears to prefer the warm summer-fall seasons, and is not targeted by the *nifH* qPCR assay for *C. watsonii*^73^. Future sampling efforts in the North Pacific Ocean are, therefore, needed to culture more *C. waterburyi* strains and quantify their environmental abundance.

#### b. *C. waterburyi* has been detected in the North Pacific Ocean across multi-year time spans

Microscopic data in **Figure 1**, shows that rod-shaped *C. waterburyi-*like cells were found in particle traps in both 2010 and 2022, suggesting that *C. waterburyi* C + N export is common and consistent in the North Pacific (and other oligotrophic ocean regions; **Figure 4D**). While these data are visually compelling, they do not immediately prove that these *Crocosphaera*-like cells are members of the clades discussed above. To test this, we read-mapped *C. waterburyi*, *C. watsonii*, and CrocoG genomes to the Stn. ALOHA 4,000 m deep trap metagenomic reads^35^ as well as other large metagenomic datasets (**Suppl. Table S7**). This effort showed that both *C. waterburyi* and *C. watsonii* were detected at >25% genome presence across all three years (2014-2016) of sampling from 4,000-meter sediment traps (**Suppl. Figure S3**) but also had different % recruitment values across these years, with *C. waterburyi* increasing in % recruitment over time (**Figure 6)** and potentially becoming more abundant in this ecosystem.

**Figure 6.**
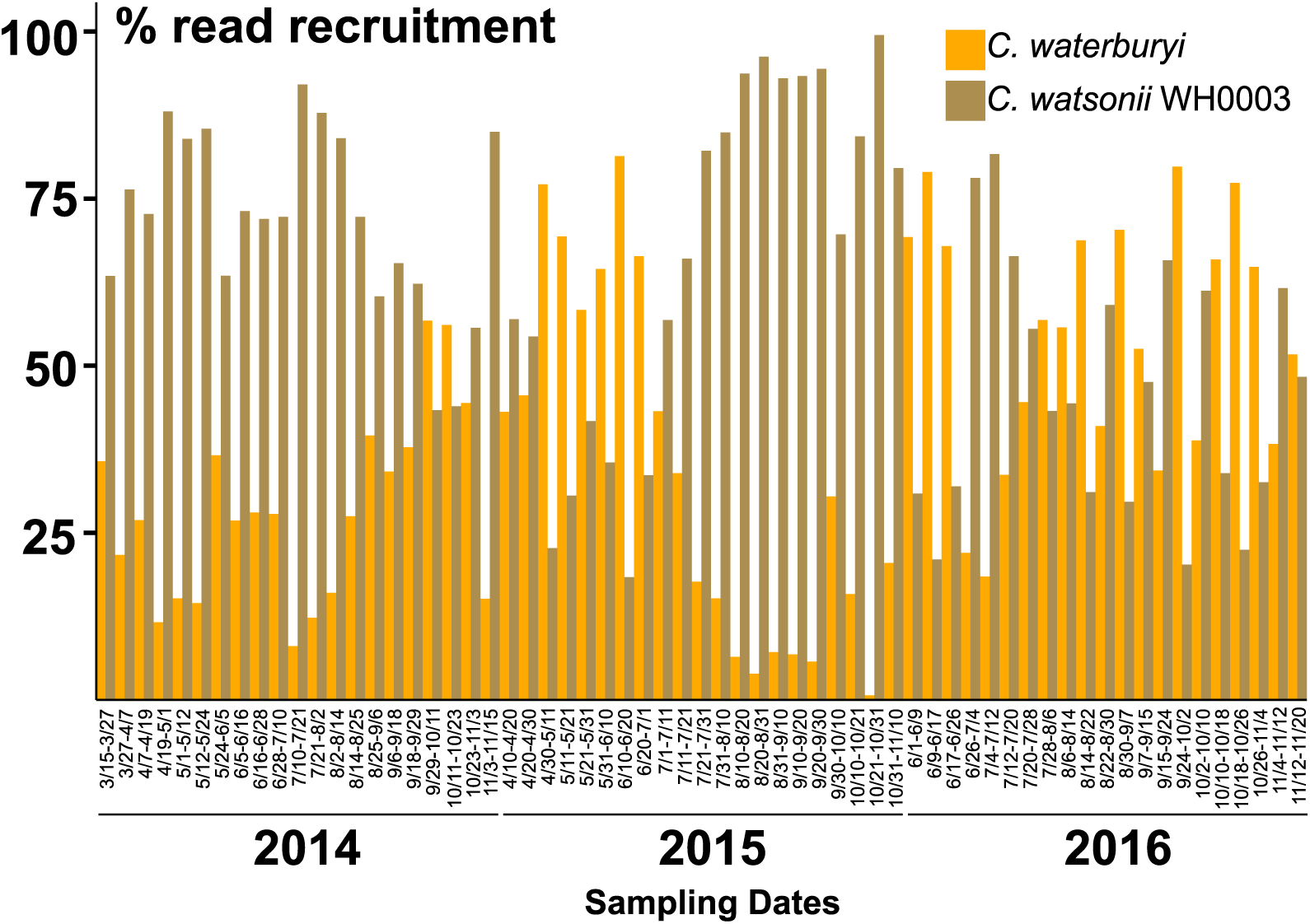
Read mapping of *C. waterburyi* and *C. watsonii* WH0003 genomes to 4,000 m sediment trap metagenomic samples from 2014-2016^35^. The % recruitment of mapped reads is shown for *C. watsonii* and *C. waterburyi* (interpretation: of the reads that were mapped, X% mapped to *C. watsonii* and X% mapped to *C. waterburyi*). The CrocoG were included in the analysis but are not shown here as their % genome detection across all samples was always <0.4%.

While *C. watsonii* was detected at a small number of stations in the BioGEOTRACES, GoShip, TaraOceans, and 150 m net trap North Pacific metagenomes (**Suppl. Table 7**), *C. waterburyi* was not detected in other large metagenomic datasets, which is likely attributed to blooms being dense yet highly ephemeral due to formation of large aggregates with faster sinking rates than *C. watsonii*. As well, the absence of *C. waterburyi* in other metagenomic datasets could be due to the difficulty of achieving cellular lysis via traditional DNA extraction kits; we were unable to extract adequate concentrations of genomic DNA with a kit until we amended the manufacturer’s protocol with 5x liquid N_2_ freeze thaws, 3x tube agitation, and an overnight (∼12 hours) proteinase K incubation (see *Materials and Methods*).

Despite DNA extraction difficulties, the presence of *C. watsonii* and *C. waterburyi* in 4,000 m sediment trap metagenomes does suggest that these organisms are present and abundant in surface waters where they originated prior to sinking out. Additionally, the CrocoG (*i.e., C. subtropica*, *C. chwakensis*, and *Cyanothece* sp. BG0011) genomes were not detected (% genome detection always <0.4%) in these deep trap samples (**Suppl. Table 6**), affirming the biogeographical differentiation within *Crocosphaera* and highlighting the ecological importance of both *C. waterburyi* and *C. watsonii* in oceanic oligotrophic systems.

*C. waterburyi* DNA in 4,000-meter sediment trap samples (**Figure 6**) indicates that the cells sank out from the euphotic zone to depth or were ingested and excreted by zooplankton grazing on EPS aggregates, as is common with *C. watsonii*^10^. Either fate has important implications for ecosystem function and trophic structure. Recently, diazotrophs have been demonstrated as underestimated players in C + N export^11^, and the discovery of *C. waterburyi* in 4,000 m sediment traps reveals that it is also influencing C + N export in a previously unknown way. Further work on *C. waterburyi* abundances and presence on sinking particles will help to tease apart C + N dynamics of this novel species; this is of particular interest as C fixation and export by photosynthetic organisms have implications for deep ocean carbon sequestration and environmental impacts.

## Conclusion

*C. waterburyi* and *C. watsonii* appear to be sympatric and active in the oligotrophic open ocean but are different species by multiple metrics (e.g., morphological, physiological, phylogenomic, pan genomic, phylogenetic, and % ANI comparisons). The characterization of *C. waterburyi* opens a new window into the ecology and distribution of unicellular N_2_-fixers in the marine environment. *Crocosphaera* are important links in the marine food web that bring new sources of organic C + N into low nutrient, oligotrophic ocean regions^4,5,9^. In a changing global climate, understanding marine microbial communities is even more essential to predicting environmental outcomes, and like other diazotrophs, single-celled *Crocosphaera* can be considered keystone species in the oligotrophic oceans^2,8^. *C. waterburyi* also grows better at high temperatures than *C. watsonii* and appears to be increasing in relative abundance in the North Pacific Ocean. Therefore, *C. waterburyi* may be a predicted climate-change warming “winner” for the oligotrophic oceans.

The discovery of *C. waterburyi* demonstrates that there is still more to be learned about oceanic N_2_-fixer diversity and highlights the need for further studies focused on the genus *Crocosphaera*, both in sinking particles and the surface ocean, to understand how they may respond and change under anthropogenic warming of the oceans.

## Data and Code Availability

The whole genome sequence of Candidatus *Crocosphaera waterburyi* Alani8 has been deposited on GenBank under BioProject PRJNA951741 and BioSample SAMN34055600. The raw forward and reverse reads are available on the NCBI Sequence Read Archive project PRJNA951741. Demultiplexed raw *nifH* amplicon sequences are available on the NCBI Sequence Read Archive under BioProject PRJNA1009239. Code for bioinformatic pipelines can be found at: https://github.com/catiecleveland/Crocosphaera-Biogeography.

## Acknowledgements

We would like to thank Lily Momper for assistance with field sampling and Anjali Bhatnagar for assistance with laboratory work. This work would not be possible without the support of the captain and crew of the R/V Kilo Moana and Chief Scientists during both the KM1013 (Chief Scientist Ben Van Mooy - WHOI) and PARAGON I cruises (Chief Scientists Angelicque White - UH Manoa, Matthew Church - Univ. Montana) and Ellen Salamon Slater and Lasse Riemann - Univ. Copenhagen) for sample collection during PARAGON I. Finally, we gratefully acknowledge Jonathan Magasin (UCSC) for their support with *nifH* amplicon processing.

## Contributions

CSC - performed all physiological experiments, bioinformatics and design for Figures 1A-B, 2A-C, 3, 4, 5D, and 6, microscopy for Figure 1A-B, and was primary manuscript writer.

EAW - collected the samples from KM1013, field microscopy for Figure 1C-D, and maintained the *C. waterburyi* Alani8 enrichment long-term (2010-present).

YZ – Designed CRISPR-cas supplemental figure S1B.

K-TK and JPZ – Sample collection for PARAGON I, collected and sequenced *nifH* amplicons, bioinformatics and design for Figures 1E and 5A-C, and particle microscopy for Figure 1E.

CSC, K-TK, YZ, JPZ, and EAW contributed to manuscript editing/writing.

