## supplemental figures & methods for "A novel, N_2_-fixing cyanobacterium present and active in the global oceans"

**Supplemental Figures S1-S3**

**Supplemental Methods and References**

**Supplemental Methods**

**CRISPR-cas Annotation**

All GenBank genomes in the phylogenomic tree (**Figure 2**) were downloaded from NCBI and annotated with CCTyper.^2^ Organisms found to have some or all Cas genes and CRISPR arrays necessary for a CRISPR-cas system were identified and superimposed on the phylogenomic tree. One CRISPR-cas system annotated as “ambiguous” was not included.

**CRISPR-cas 8b Gene Phylogenetics**

Nine organisms in the family *Aphanothecacea*, when annotated by CCTyper,^2^ were shown to encode a Type I-B CRISPR-cas system. For each of these nine, a contigs database was generated using Anvi’o v7.1^3^. The contigs databases were visualized and subsequently the Cas8b diagnostic gene for the Type I-B system^4^ were found, identified, and the sequences were exported for alignment. The sequences were aligned using Clustal Omega 1.2.2^5^, trimmed, and a Cas8b gene tree was created using RAxML 8.2.11^6^ with a GTR GAMMA nucleotide model, rapid bootstrapping (1000 bootstraps), and a search for best-scoring maximum likelihood tree algorithm.

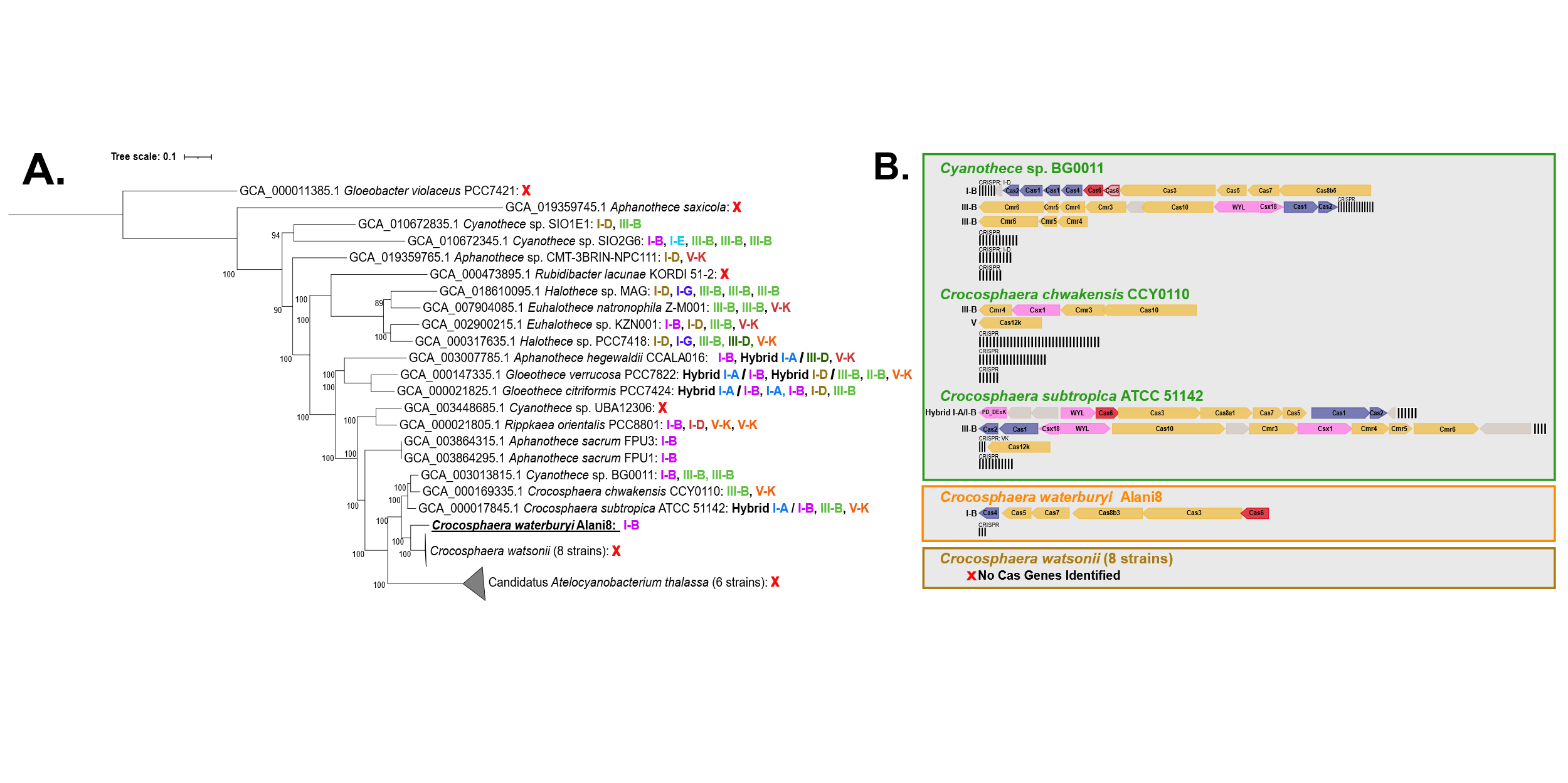

**S1.** CRISPR-cas annotation of 35 unicellular N_2_-fixing cyanobacteria **(A)** and the specific cas genes and CRISPR arrays in genus *Crocosphaera* **(B)**. *C. waterburyi* and the CrocoG encode cas genes, CRISPR arrays, and spacer sequences. In **(B)**, yellow shows the interference module, blue shows the adaptation module, purple shows the accessory genes, and red shows Cas6.

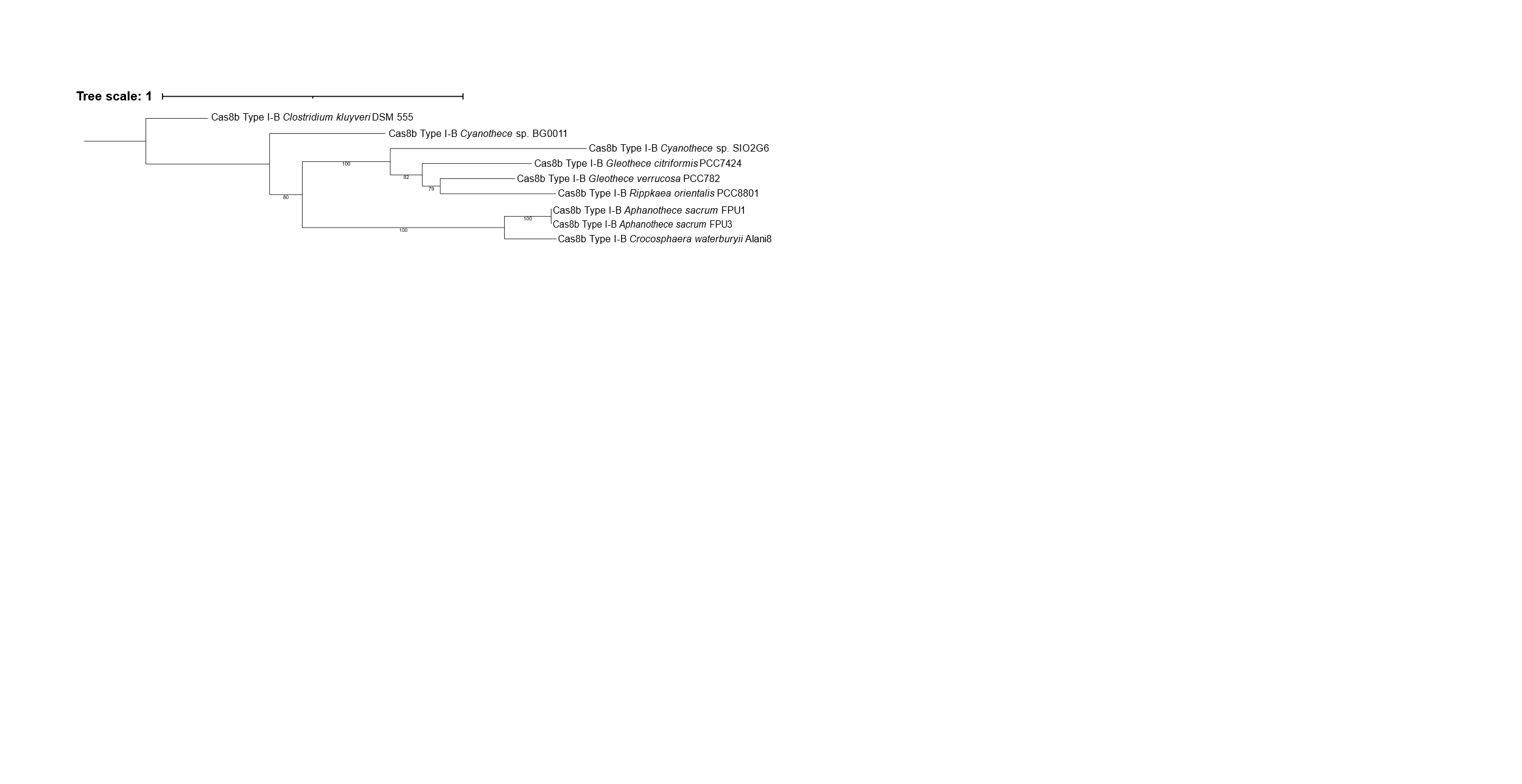

**S2.** The Cas8b DNA sequences (diagnostic gene for Type I-B system) from the organisms in the family *Aphanothecacea* that encoded a Type I-B CRISPR-cas system compiled into a phylogenetic tree. Only bootstraps above 75 are shown.

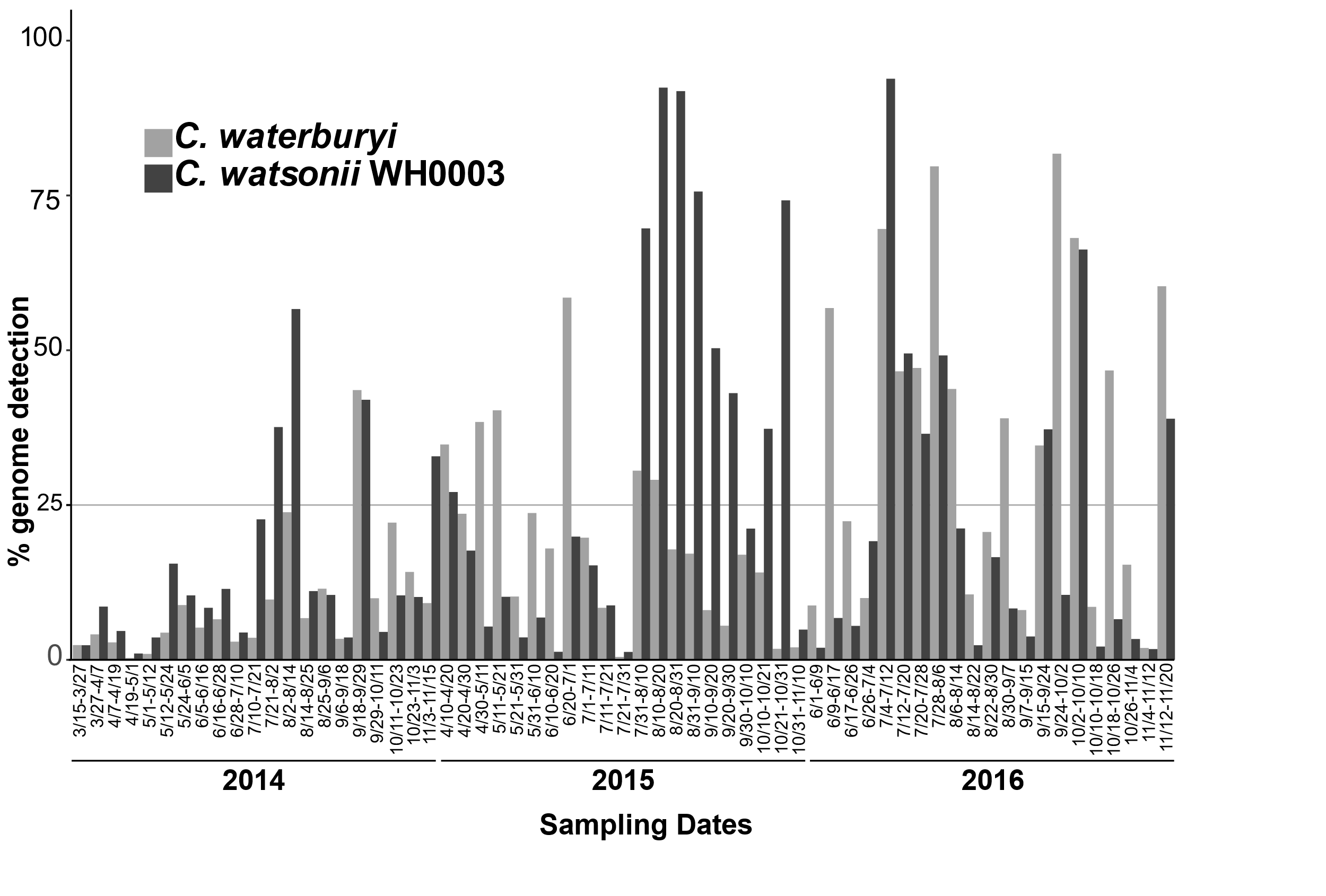

**S3.** The % genome detection of *C. waterburyi* and *C. watsonii* is shown across the deep trap metagenomes^1^ from 2014-2016. Both *C. watsonii* and *C. waterburyi* were at >25% genome detection across all 3 years in the North Pacific Ocean, indicating that they are consistently present in this environment. The CrocoG were included in the analysis but were not significantly detected thus are not shown here.
